# Impact of index hopping and bias towards the reference allele on accuracy of genotype calls from low-coverage sequencing

**DOI:** 10.1101/358085

**Authors:** Roger Ros-Freixedes, Battagin Mara, Martin Johnsson, Gregor Gorjanc, Alan J Mileham, Steve D Rounsley, John M Hickey

## Abstract

**Background:** Inherent sources of error and bias that affect the quality of the sequence data include index hopping and bias towards the reference allele. The impact of these artefacts is likely greater for low-coverage data than for high-coverage data because low-coverage data has scant information and standard tools for processing sequence data were designed for high-coverage data. With the proliferation of cost-effective low-coverage sequencing there is a need to understand the impact of these errors and bias on resulting genotype calls.

**Results:** We used a dataset of 26 pigs sequenced both at 2x with multiplexing and at 30x without multiplexing to show that index hopping and bias towards the reference allele due to alignment had little impact on genotype calls. However, pruning of alternative haplotypes supported by a number of reads below a predefined threshold, a default and desired step for removing potential sequencing errors in high-coverage data, introduced an unexpected bias towards the reference allele when applied to low-coverage data. This bias reduced best-guess genotype concordance of low-coverage sequence data by 19.0 absolute percentage points.

**Conclusions:** We propose a simple pipeline to correct this bias and we recommend that users of low-coverage sequencing be wary of unexpected biases produced by tools designed for high-coverage sequencing.

## Introduction

Sequence data has the potential to empower identification of causal variants underlying quantitative traits or diseases, to enhance livestock breeding, and to increase the precision and scope of population genetic studies. For sequence data to be used routinely in research and breeding, low-cost sequencing strategies must be used to assemble large data sets covering most of the genetic diversity in a population. Such low-cost strategies could involve sequencing individuals at low coverage followed by imputation [1–3].

Current sequencing technologies have inherent sources of errors and bias that affect the quality of the sequence data [4–6]. Two of the most important are index hopping and bias towards the reference allele. The impact of these artefacts is likely greater for low-coverage data than for high-coverage data because low-coverage data has scant information and standard tools for processing sequence data were designed for high-coverage data. With the proliferation of cost-effective low-coverage sequencing there is a need to understand the impact of these artefacts on resulting genotype calls.

Index hopping has a biochemical cause and appears in the early stages of sequencing. Currently, the most widely used high-throughput sequencing platform is the HiSeq series of instruments from Illumina Inc. Due to the large sequencing capacity of these platforms, several samples are often sequenced jointly within a single flow cell channel by multiplexing. To link multiplexed sequence reads to the original samples, the adapter sequences used during library preparation include a set of unique index sequences. However, molecular recombination of indices, or ‘index hopping’, can occur when free adapters are present in a library pool. This leads to misassignment of sequence reads between samples in the multiplex. Recently alarming data showed index hopping incidences of up to 10% [7]. These results sparked debate and concern about index hopping, though some subsequent studies reported a low incidence for most applications [8–10], which is in line with expectation provided that cleaning protocols are used to remove free adapters from the libraries [11]. While these results are reassuring, they pertained to high-coverage sequence data and the effect of index hopping on low-coverage sequence data and its downstream analysis remains unclear.

Bias towards the reference allele can be observed in sequence data following bioinformatic processing. It originates mainly during read alignment, but it can also occur during variant discovery and genotyping. Alignment of sequence reads onto a haploid reference genome relies on the calculation of similarity scores between reads and the reference genome. The more a read diverges from the reference, the more unlikely it is to align appropriately. This disfavours the alignment of reads that carry the alternative allele at a variant position because such reads have at least one more mismatch to the reference genome compared to reads that carry the reference allele. If a read covers multiple variant sites and carries alternative alleles at multiple positions, the probability of aligning such a read decreases even further, which in turn produces a stronger reference allele bias in highly polymorphic regions. This can lead to biases in downstream applications, e.g., in estimation of allele frequencies [4,12].

Another potential source of bias towards the reference allele can occur during variant discovery and genotyping. One of the most popular variant callers is GATK HaplotypeCaller [13], which provides a pipeline for efficient joint genotyping of multiple samples. In the GATK Best Practices pipeline, variant discovery and joint genotyping of multiple samples are performed as two separate steps [13,14]. In the variant discovery step, read information for each position of the reference genome is stored for each individual sample in a gVCF file, which differs from the traditional VCF file in that it stores information of the non-variant positions as well as the variant positions. In the joint genotyping step, the gVCF files that have been created separately for each individual are combined and genotypes are called for all individuals at all the positions that are variant for at least one individual in the sequenced population. Compared to other pipelines, this two-step process has the advantage that only the genotyping (and not the variant calling itself, which is the most computationally demanding step) is done jointly for all the samples. This improves scalability and facilitates the incorporation of new batches of sequenced individuals for the joint genotyping step. However, GATK HaplotypeCaller was designed for high-coverage sequencing and, to our knowledge, its performance in low-coverage sequencing has not been assessed.

In this study we explored the impact of index hopping and bias towards the reference allele in low-coverage sequencing. We show that index hopping and bias towards the reference allele due to alignment have little impact on genotype calls. However, unexpected biases may arise from pipelines that use tools designed for high-coverage sequence data when applied to low-coverage sequence data. In particular we describe how a function from GATK HaplotypeCaller that is very useful for high-coverage data introduces a strong bias towards the reference allele when used on low-coverage data. We propose a new pipeline that avoids this bias. The results in this paper show the importance of validating the performance of tools designed for high-coverage data on low-coverage data.

## Materials and Methods

### Sequenced individuals

A total of 26 commercial pigs were used in this study. Tissue samples were collected from ear punches or tail clippings and genomic DNA was extracted using Qiagen DNeasy 96 Blood & Tissue kits (Qiagen Ltd., Mississauga, ON, Canada). Paired-end library preparation was conducted using the TruSeq DNA PCR-free protocol (Illumina, San Diego, CA). Two sets of libraries were produced; one with average insert size of 350 bp and the other with average insert size of 550 bp. Libraries with average insert size of 350 bp were sequenced on a HiSeq 4000 instrument, for a target coverage of 2x per sample. For this, all 26 samples were multiplexed within a single flow cell channel. Libraries with average insert size of 550 bp were sequenced on a HiSeq X instrument, for a target coverage of 30x per sample. For this, the 26 samples were sequenced one sample per flow cell channel. All libraries were sequenced at Edinburgh Genomics (Edinburgh Genomics, University of Edinburgh, Edinburgh, UK). DNA samples from the same pigs were also genotyped using the GGP-Porcine HD BeadChip (GeneSeek, Lincoln, NE).

### Variant discovery

DNA sequence reads were pre-processed using Trimmomatic [15] to remove adapter sequences from the reads. The reads were then aligned to the reference genome *Sscrofa11.1* (GenBank accession: GCA_000003025.6) using the BWA-MEM algorithm [16]. Duplicates were marked with Picard (http://broadinstitute.github.io/picard). Single nucleotide polymorphisms (SNPs) and short insertions and deletions (indels) were identified with the variant caller GATK HaplotypeCaller (GATK 3.8.0; [13,14]). The GATK HaplotypeCaller performs local re-assembly of the reads to generate a list of possible haplotypes in a region by constructing a read-threading graph. Sections of that graph that are supported by a number of reads (kmers) lower than a predefined threshold are considered likely to be sequencing errors and removed from the graph in a step referred to as ‘pruning’. By default the threshold for pruning is set to ‘–minPruning 2’. We used the default settings but we also performed variant discovery without pruning (–minPruning 1). Variant discovery with GATK HaplotypeCaller was performed separately for each individual. A joint variant set for the 26 individuals was obtained by extracting the variant positions from all the individuals with GATK GenotypeGVCFs. Finally, variants were filtered with VCFtools [17] and only biallelic SNPs were retained. To minimise computing costs, we only considered variants on chromosome 1.

### Genotyping

We did not use genotypes called directly by GATK GenotypeGVCFs or any software tool. Instead we extracted allele read counts (i.e., the coverage that each allele received at each variant site) from the VCF file. We then called genotypes based on genotype probabilities calculated from allele read counts of the reference allele (nRef) and the alternative allele (nAlt). Genotype probabilities for the reference homozygote (0), heterozygote (1), and alternative homozygote (2) were respectively calculated as:

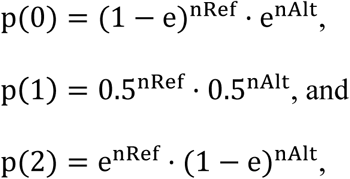

where e is the sequencing error rate, assumed to be 0.01. The three probabilities were scaled to sum up to 1. Genotype calls were made with three different levels of certainty: (i) the most probable genotypes (referred to as ‘best-guess’); (ii) genotypes that had a probability greater than 0.90; or (iii) genotypes that had a probability greater than 0.98.

### Genotype and allele concordance

Genotype concordance was calculated by: (i) comparing genotypes for the same variant from the sequence data and the marker array, using the marker array genotypes as the true genotypes; or (ii) comparing the same variant from the sequence data at low and high coverage and using the high-coverage genotype calls as the true genotypes. Genotype concordance was calculated as the percentage of matches between the nominal true genotypes and the genotype calls. We only considered genotypes for SNPs that segregated in the marker array and in the sequence data. The number of SNPs tested for concordance with marker array data was 5,136 for the low-coverage data and 5,531 for the high-coverage data. The set of 5,531 SNPs was used for testing the concordance between low- and high-coverage sequence data. We also calculated allele concordance, as the percentage of matched alleles between the nominal true genotypes and the genotype calls.

### Bias towards the reference allele due to variant caller and new pipeline

Initially we called genotypes using the read counts stored in the gVCF files produced by GATK HaplotypeCaller. For the testing potential biases introduced by the variant caller, we also called genotypes using the read counts obtained directly from the aligned reads stored in the BAM files. To do so, we extracted the read counts from the BAM files for variant sites discovered using pysam (https://github.com/pysam-developers/pysam). We refer to this method as the ‘new’ pipeline.

Our initial results indicated that there was a strong bias towards the reference allele introduced by the variant caller. Therefore, for all further analyses we used read counts obtained from the BAM files with the new pipeline for genotyping. We called genotypes for the 5,531 variant positions on chromosome 1 discovered from the high-coverage sequence data that had already been genotyped using the marker array.

### Bias towards the reference allele due to alignment

In this study we defined alignment bias to be the differential alignment of almost-identical reads that differ only in one allele at a given variant position, be it either the reference allele or the alternative allele. To quantify the alignment bias, we aligned the 2x data against two reference genomes: the ‘original’ reference genome and a ‘tailored’ reference genome. The tailored reference genome was created by replacing the reference allele with the alternative allele at all the variant positions discovered with the 30x sequence data in chromosome 1. Thus, the allele that was originally the alternative allele became the reference allele in the tailored reference genome and vice versa. We extracted the allele read counts from the aligned reads in the BAM files generated with both reference genomes. The allele read counts were used to call genotypes for evaluating the genotype concordance between the 2x data and the true genotypes (from the 30x data). Genotypes were called from the allele read counts obtained with either: (i) the original reference genome (**REF**), or (ii) the tailored reference genome (**ALT**). Because REF could favour the alignment of reads carrying the reference allele and disfavour the alignment of reads carrying the alternative allele, and vice versa for ALT, we also considered two additional cases that were a combination of the previous two: (iii) read counts for the reference allele from the original reference genome and read counts for the alternative allele from the tailored reference genome (**CIS**), and (iv) vice versa, read counts for the reference allele from the tailored reference genome and read counts for the alternative allele from the original reference genome (**TRANS**). Thus, the CIS case used allele read counts that had more favourable alignment for each allele, and, on the contrary, the TRANS case used allele read counts that had more unfavourable alignment for each allele.

### Index hopping

In order to quantify the incidence of index hopping in our 2x dataset, we generated 2x data that were either free of index hopping or had different levels of index hopping simulated. The 2x data free of index hopping were generated by downsampling the 30x data (i.e., random sampling of ~1/15 of the 30x reads), which had been generated without multiplexing (1 sample per lane). The downsampled 2x data was used to obtain baseline sequence data in the absence of index hopping. We then added index hopping to this data by deliberately assigning reads to other individuals at random with a probability of 0.1%, 0.5%, 1%, 2%, or 5%. For each of these cases we downsampled the data independently before simulating index hopping to account for the random sampling of reads that occurs during sequencing.

To analyse the data, genotypes in each dataset were called as we described above (best-guess or above a certain probability threshold) but also with an additional method based on the presence/absence of each allele that was more sensitive to index hopping. With this presence/absence method, the presence of a single read supporting the opposite allele was sufficient to change the genotype call (e.g., the genotype call with nRef=10 and nAlt=0 would be homozygote but the genotype call with nRef=10 and nAlt=1 would be heterozygote). Note that this method is equivalent to calling best-guess genotypes with null sequencing error rate.

We regressed the percentages of genotype concordance on the level of index hopping and used this regression to predict the level of index hopping level in the observed 2x dataset. Concordance percentages represent relative, rather than absolute, information and therefore should not be analysed using standard statistical techniques that are defined in real space, which has an absolute scale [18]. In order to validate the results with a methodology that was more appropriate for compositional data we also analysed the data using isometric log-ratio transformations (**ilr**) of the concordance percentages [19,20]. The ilr were the log-ratios of the percentage of correct calls against the percentages of incorrect calls or the log-ratios of the percentage of correct homozygous calls against the percentage of incorrect heterozygous calls. For the ilr variables a quadratic regression was fitted.

## Results

### Variant discovery

Most of the SNPs present on the marker array were discovered using sequence data, either at high or low coverage. The number of biallelic SNPs discovered on chromosome 1 with high- and low-coverage data is shown in Table 1. A total of 1,693,308 biallelic SNPs were discovered with the high-coverage data and 1,333,943 with the low-coverage data. The low-coverage sequence data contained 76.3% (1,292,269) of the biallelic SNPs discovered with the high-coverage data. The marker array had 5,779 SNPs on chromosome 1 that segregated in the 26 sequenced individuals. Of these, 95.7% (5,531) were discovered with the high-coverage data and 88.9% (5,136) with the low-coverage data.

**Table 1.**
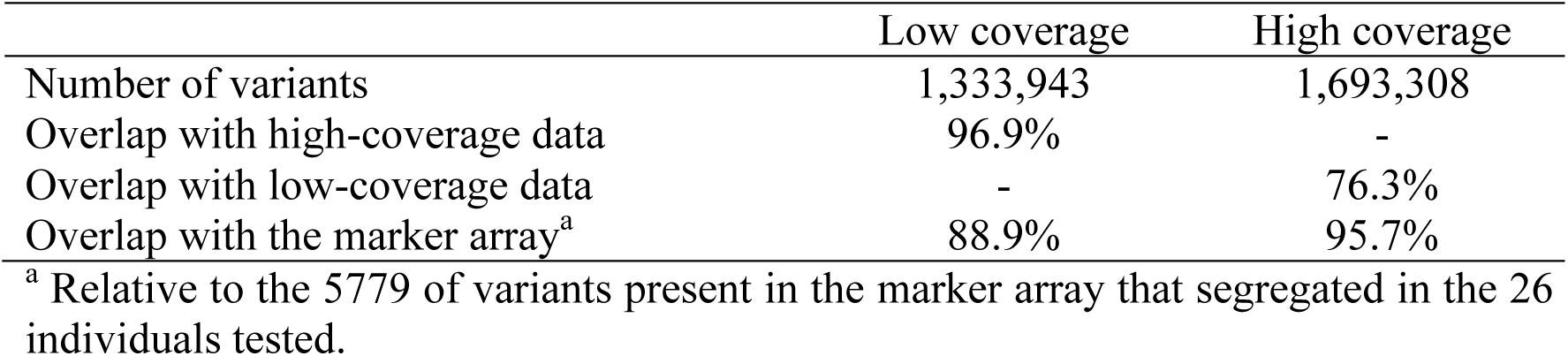
Number of biallelic SNPs discovered in chromosome 1 at low and high coverage and overlap with the marker array.

|  | Low coverage | High coverage |
| --- | --- | --- |
| Number of variants | 1,333,943 | 1,693,308 |
| Overlap with high-coverage data | 96.9% | - |
| Overlap with low-coverage data | - | 76.3% |
| Overlap with the marker array<sup>a</sup> | 88.9% | 95.7% |

<sup>a</sup> Relative to the 5779 of variants present in the marker array that segregated in the 26 individuals tested.

Disabling the pruning step in GATK HaplotypeCaller for processing the low-coverage data increased the number of variants discovered but also the number of potential false positives. The number of biallelic SNPs discovered on chromosome 1 with low-coverage data with or without pruning is shown in Table 2. When pruning was disabled, a total of 1,877,644 biallelic SNPs were discovered with the low-coverage data. This number was greater than the set of variants discovered with the high-coverage data with the default pruning settings (Table 1). However, 24.1% of those extra SNPs could not be validated using the high-coverage data, which is a much greater proportion than when pruning was used (3.1%).

**Table 2.** Number of biallelic SNPs discovered in chromosome 1 at low coverage with different GATK HaplotypeCaller pruning options along with percentage not validated at high coverage and genotype and allele concordances with the marker array.

|  | minPruning=2<br>(default) | minPruning=1 |
| --- | --- | --- |
| Number of variants | 1,333,943 | 1,877,644 |
| Not validated at high coverage | 3.1% | 24.1% |
| Best-guess genotype concordance | 62.1% | 76.5% |
| Allele concordance | 77.6% | 87.5% |

### Genotype concordance and bias towards reference allele due to variant calling

The variant caller that we used introduced a bias towards the reference allele and this had a great impact on genotype calling with low-coverage data. Table 3 shows the genotype concordance for calls obtained with the allele read counts from the gVCF files produced by GATK HaplotypeCaller. The table shows a large bias towards the reference allele for low-coverage sequence data. In the most extreme case of positions with 1x, we would expect the genotypes that are heterozygous according to the marker array to be called as either one of the two possible homozygotes ‘0’ and ‘2’ 50% of the times. Instead we called them as the reference homozygote ‘0’ 95.1% of the times and as the alternative homozygote ‘2’ only 4.9% of the times. Also, at 1x, 82.0% of the alternative homozygotes ‘2’ were called as reference homozygote ‘0’. Because of this bias, the overall genotype concordance was only 62.1% and the allele concordance was only 77.6%.

**Table 3.**
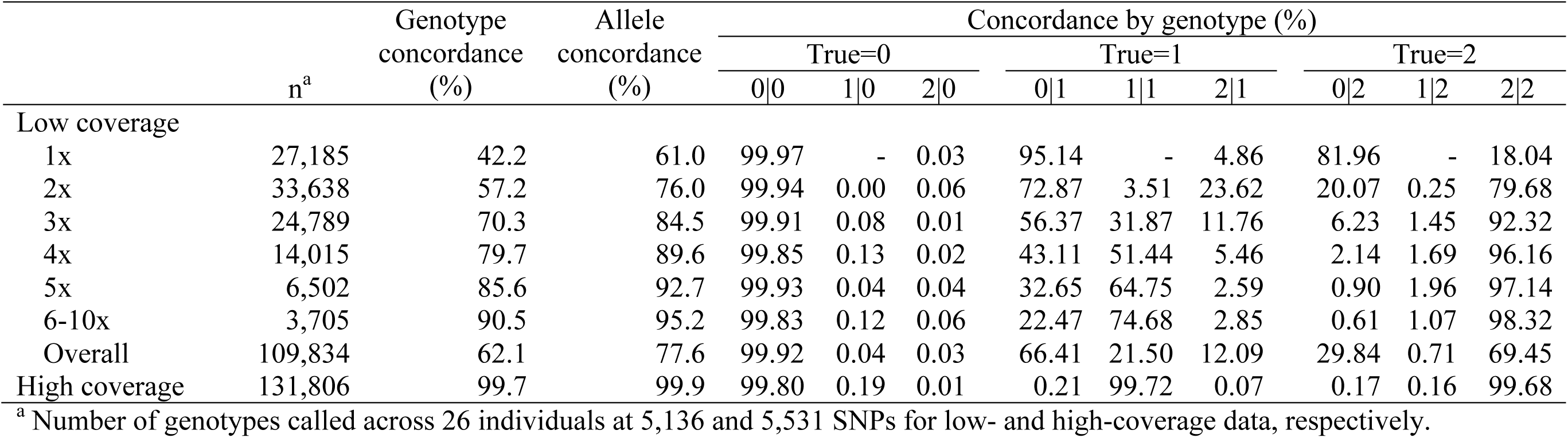
Concordance between best-guess genotype calls from sequence data and marker array genotypes with the allele read counts obtained with the default settings of GATK HaplotypeCaller. Concordance is shown by coverage at variant position.

|  | n<sup>a</sup> | Genotype<br>concordance<br>(%) | Allele<br>concordance<br>(%) | Concordance by genotype (%) |  |  |  |  |  |  |  |  |
| --- | --- | --- | --- | --- | --- | --- | --- | --- | --- | --- | --- | --- |
|  |  |  |  | True=0 |  |  | True=1 |  |  | True=2 |  |  |
|  |  |  |  | 0\|0 | 1\|0 | 2\|0 | 0\|1 | 1\|1 | 2\|1 | 0\|2 | 1\|2 | 2\|2 |
| Low coverage |  |  |  |  |  |  |  |  |  |  |  |  |
| 1x | 27,185 | 42.2 | 61.0 | 99.97 | - | 0.03 | 95.14 | - | 4.86 | 81.96 | - | 18.04 |
| 2x | 33,638 | 57.2 | 76.0 | 99.94 | 0.00 | 0.06 | 72.87 | 3.51 | 23.62 | 20.07 | 0.25 | 79.68 |
| 3x | 24,789 | 70.3 | 84.5 | 99.91 | 0.08 | 0.01 | 56.37 | 31.87 | 11.76 | 6.23 | 1.45 | 92.32 |
| 4x | 14,015 | 79.7 | 89.6 | 99.85 | 0.13 | 0.02 | 43.11 | 51.44 | 5.46 | 2.14 | 1.69 | 96.16 |
| 5x | 6,502 | 85.6 | 92.7 | 99.93 | 0.04 | 0.04 | 32.65 | 64.75 | 2.59 | 0.90 | 1.96 | 97.14 |
| 6-10x | 3,705 | 90.5 | 95.2 | 99.83 | 0.12 | 0.06 | 22.47 | 74.68 | 2.85 | 0.61 | 1.07 | 98.32 |
| Overall | 109,834 | 62.1 | 77.6 | 99.92 | 0.04 | 0.03 | 66.41 | 21.50 | 12.09 | 29.84 | 0.71 | 69.45 |
| High coverage | 131,806 | 99.7 | 99.9 | 99.80 | 0.19 | 0.01 | 0.21 | 99.72 | 0.07 | 0.17 | 0.16 | 99.68 |

<sup>a</sup> Number of genotypes called across 26 individuals at 5,136 and 5,531 SNPs for low- and high-coverage data, respectively.

The bias towards the reference allele due to variant caller can be avoided by calling genotypes from the read counts obtained directly from the aligned reads stored in BAM files. Table 4 shows the genotype concordance obtained with the new pipeline using allele read counts extracted directly from BAM files. The bias was corrected and the concordances matched expectations. Overall, genotype and allele concordances rose to 81.1% and 90.5%, respectively. As expected, most of the incorrect calls arose from the difficulty of calling heterozygous genotypes at low coverage.

**Table 4.** Concordance between best-guess genotype calls from sequence data and marker array genotypes with the allele read counts obtained from the aligned reads in the BAM files. Concordance is shown by coverage at variant position.

|  | n<sup>a</sup> | Genotype<br>concordance<br>(%) | Allele<br>concordance<br>(%) | Concordance by genotype (%) |  |  |  |  |  |  |  |  |
| --- | --- | --- | --- | --- | --- | --- | --- | --- | --- | --- | --- | --- |
|  |  |  |  | True=0 |  |  | True=1 |  |  | True=2 |  |  |
|  |  |  |  | 0\|0 | 1\|0 | 2\|0 | 0\|1 | 1\|1 | 2\|1 | 0\|2 | 1\|2 | 2\|2 |
| Low coverage |  |  |  |  |  |  |  |  |  |  |  |  |
| 1x | 28,300 | 62.1 | 80.8 | 99.34 | - | 0.66 | 51.46 | - | 48.54 | 0.96 | - | 99.04 |
| 2x | 32,699 | 79.5 | 89.7 | 98.42 | 1.53 | 0.05 | 26.41 | 48.15 | 25.44 | 0.21 | 1.70 | 98.09 |
| 3x | 25,993 | 88.3 | 94.1 | 98.25 | 1.72 | 0.03 | 14.01 | 71.98 | 14.01 | 0.12 | 2.36 | 97.52 |
| 4x | 16,346 | 92.5 | 96.3 | 97.91 | 2.09 | 0.00 | 8.36 | 83.84 | 7.80 | 0.00 | 2.77 | 97.23 |
| 5x | 8,878 | 94.9 | 97.5 | 97.28 | 2.72 | 0.00 | 4.83 | 91.15 | 4.02 | 0.16 | 2.81 | 97.03 |
| 6-10x | 6,444 | 95.0 | 97.5 | 97.43 | 2.50 | 0.07 | 5.01 | 91.09 | 3.90 | 0.00 | 2.75 | 97.25 |
| Overall | 118,660 | 81.1 | 90.5 | 98.39 | 1.43 | 0.18 | 24.43 | 52.30 | 23.27 | 0.33 | 1.71 | 97.96 |
| High coverage | 131,782 | 99.8 | 99.9 | 99.80 | 0.19 | 0.01 | 0.12 | 99.81 | 0.07 | 0.10 | 0.17 | 99.73 |

Disabling pruning was not as good a solution for correcting the bias as the new pipeline of extracting the allele read counts from the BAM files. Table 2 shows genotype and allele concordances with the default pruning setting and without pruning. Without pruning, the genotype and allele concordances rose to 76.5% and 87.5%, respectively, but these percentages were lower than with the new pipeline.

Once the bias towards the reference allele due to the variant caller was corrected, the concordance at homozygous positions was very high regardless of the conservativeness of the genotype calls, but these thresholds were important for concordance at heterozygous positions. Table 5 shows genotype concordance between calls with low- and high-coverage data obtained as best-guess genotypes or with a minimum probability of 0.90 or 0.98. At homozygous positions, the best-guess genotypes had an overall concordance of 98.5% and 98.2%, which was greater than the concordance of the calls with a minimum probability of 0.90 (97.2% and 96.4%, respectively), despite the latter being called with a greater level of certainty. The reason for this is that with a minimum probability of 0.90, there is not enough certainty for calling any genotype at 1x, and at positions with coverage of 2x or 3x only potential heterozygotes (either true or false), but not homozygotes, can be called due to the considered error rate. While the number of homozygotes incorrectly called as heterozygous was actually very low, the impact of these incorrect calls on the overall concordance was noticeable because the low-coverage data had many more loci with 2x and 3x than with 4x or more. A similar situation happened with genotype calls with a minimum probability of 0.98.

**Table 5.** Concordance between genotype calls with different levels of conservativeness from low- and high-coverage sequence data with the allele read counts obtained from the aligned reads in the BAM files. Concordance is shown by coverage at variant position.

|  | n<sup>a</sup> | Genotype<br>concordance<br>(%) | Allele<br>concordance<br>(%) | Concordance by genotype (%) |  |  |  |  |  |  |  |  |
| --- | --- | --- | --- | --- | --- | --- | --- | --- | --- | --- | --- | --- |
|  |  |  |  | True=0 |  |  | True=1 |  |  | True=2 |  |  |
|  |  |  |  | 0\|0<sup>b</sup> | 1\|0 | 2\|0 | 0\|1 | 1\|1 | 2\|1 | 0\|2 | 1\|2 | 2\|2 |
| Best-guess |  |  |  |  |  |  |  |  |  |  |  |  |
| 1x | 30,875 | 62.6 | 81.1 | 99.45 | - | 0.55 | 51.35 | - | 48.65 | 0.70 | - | 99.30 |
| 2x | 35,688 | 79.9 | 90.0 | 98.60 | 1.40 | 0.00 | 26.20 | 48.26 | 25.54 | 0.04 | 1.61 | 98.35 |
| 3x | 28,357 | 88.6 | 94.3 | 98.36 | 1.64 | 0.00 | 13.80 | 72.20 | 14.00 | 0.00 | 2.22 | 97.78 |
| 4x | 17,849 | 92.7 | 96.4 | 98.05 | 1.95 | 0.00 | 8.22 | 84.01 | 7.77 | 0.00 | 2.63 | 97.37 |
| 5x | 9,619 | 95.3 | 97.6 | 97.40 | 2.60 | 0.00 | 4.55 | 91.52 | 3.93 | 0.00 | 2.49 | 97.51 |
| 6-10x | 7,047 | 95.2 | 97.6 | 97.73 | 2.27 | 0.00 | 4.96 | 91.05 | 4.00 | 0.00 | 2.68 | 97.32 |
| Overall | 129,435 | 81.4 | 90.7 | 98.53 | 1.34 | 0.13 | 24.27 | 52.40 | 23.33 | 0.18 | 1.61 | 98.21 |
| Probability ≥ 0.90 |  |  |  |  |  |  |  |  |  |  |  |  |
| 1x | 0 | - | - | - | - | - | - | - | - | - | - | - |
| 2-3x<sup>b</sup> | 14,572 | 95.5 | 97.7 | - | 100.00 | - | - | 100.00 | - | - | 100.00 | - |
| 4x | 14,359 | 92.7 | 96.3 | 99.97 | 0.03 | 0.00 | 16.24 | 68.39 | 15.37 | 0.00 | 0.05 | 99.95 |
| 5x | 8,315 | 96.3 | 98.2 | 99.92 | 0.08 | 0.00 | 6.76 | 87.41 | 5.83 | 0.00 | 0.10 | 99.90 |
| 6-10x | 6,397 | 98.1 | 99.0 | 99.83 | 0.17 | 0.00 | 3.08 | 94.65 | 2.27 | 0.00 | 0.29 | 99.71 |
| Overall | 43,643 | 95.1 | 97.5 | 97.18 | 2.82 | 0.00 | 3.52 | 93.26 | 3.21 | 0.00 | 3.65 | 96.35 |
| Probability ≥ 0.98 |  |  |  |  |  |  |  |  |  |  |  |  |
| 1-3x | 0 | - | - | - | - | - | - | - | - | - | - | - |
| 4-5x<sup>b</sup> | 4,366 | 99.8 | 99.9 | - | 100.00 | - | - | 100.00 | - | - | 100.00 | - |
| 6-10x | 6,313 | 98.1 | 99.1 | 99.83 | 0.17 | 0.00 | 3.15 | 94.58 | 2.26 | 0.00 | 0.29 | 99.71 |
| Overall | 10,679 | 98.8 | 99.4 | 99.65 | 0.35 | 0.00 | 1.00 | 98.28 | 0.72 | 0.00 | 0.57 | 99.43 |

<sup>a</sup> Number of genotypes called for 5,531 SNPs across 26 individuals.
<sup>b</sup> Heterozygotes are easier to call than homozygotes. At these coverages there is not enough certainty to call the homozygotes. Note, however, that the actual counts for (1|0) and (1|2) are very low compared to (1|1): 19-fold and 569-fold lower for genotypes called with a probability greater than 0.90 and 0.98, respectively.

At heterozygous loci, it was very difficult to call heterozygotes at the lowest coverages. Because of the large proportion of loci with low coverage, the genotype concordance of heterozygous loci with best-guess genotypes was 52.4%. With more conservative calls the heterozygotes were called more accurately and the genotype concordance was 93.3% and 98.3% respectively with a minimum probability of 0.90 and 0.98. However, there was a trade-off between the concordance of called genotypes and the number of called genotypes. With more conservative calls, the number of called genotypes was only 33.7%, with a minimum probability of 0.90, or 8.3%, with a minimum probability of 0.98, of those that could be called using best-guess genotypes.

### Bias towards reference allele due to alignment

Reads with an allele that was present in the reference genome had a greater probability of successful alignment, but the difference was small. Table 6 shows the average allele read counts depending on which allele was in the reference genome. Approximately 1.3% of reads were not aligned when the reference genome contained the opposite allele than the read. The alignment bias also caused that the number of reads that carried the allele in the reference genome but were incorrectly mapped to a position where the individual was homozygous for the opposite allele increased by 9.8%, although these potentially mismapped reads represented only a small fraction of the total.

**Table 6.** Average allele read counts depending on which allele is in the reference genome.

| Allele<sup>a</sup> | Reference genome | Allele in reference genome<sup>a</sup> | Overall | True genotype |  |  |
| --- | --- | --- | --- | --- | --- | --- |
|  |  |  |  | 0 | 1 | 2 |
| Reference | Original | Reference | 1.483 | 2.470 | 1.237 | 0.017 |
|  | Tailored | Alternative | 1.463 | 2.440 | 1.219 | 0.016 |
|  |  | *Difference not aligned* | *1.3%* | *1.2%* | *1.5%* | *9.8%* |
| Alternative | Original | Reference | 0.980 | 0.014 | 1.217 | 2.407 |
|  | Tailored | Alternative | 0.993 | 0.016 | 1.234 | 2.438 |
|  |  | *Difference not aligned* | *1.3%* | *9.8%* | *1.3%* | *1.3%* |

<sup>a</sup> Alleles are defined as reference or alternative referring to the original pig reference genome *Sscrofa11.1* (GenBank assembly accession: GCA\_000003025.6).

However, the impact of the bias towards reference allele due to alignment on the genotype calls is likely to be low. Table 7 shows the genotype concordance between low- and high-coverage sequence data after alignment with the original reference genome (REF), the tailored reference genome (ALT), or a combination of both (CIS and TRANS). Using the REF or ALT reference genomes introduced some bias towards homozygotes calls for the reference or the alternative allele, respectively. Using the CIS combination, where the allele read counts were obtained from the most favourable case for each of them (i.e., the reference genome contained that same allele), increased the number of genotype calls regardless of the conservativeness of the calls and it increased the ability of correctly call heterozygotes with lower levels of certainty. On the contrary, using the TRANS combination, where the allele read counts were obtained from the least favourable case for each of them (i.e., the reference genome contained the opposite allele), reduced the number of genotype calls and the ability to correctly call heterozygotes. Overall, changes in best-guess genotype concordance were small and the percentage of incorrect calls between CIS (most favourable case) and REF (current practice) differed only by 0.1 absolute percentage points.

**Table 7.** Impact of bias towards reference allele due to alignment on concordance between low- and high-coverage sequence data by alignment with the original reference genome (REF), the tailored reference genome (ALT), or a combination of both (CIS and TRANS).

|  | n<sup>a</sup> | Genotype<br>concordance<br>(%) | Allele<br>concordance<br>(%) | Concordance by genotype (%) |  |  |  |  |  |  |  |  |
| --- | --- | --- | --- | --- | --- | --- | --- | --- | --- | --- | --- | --- |
|  |  |  |  | True=0 |  |  | True=1 |  |  | True=2 |  |  |
|  |  |  |  | 0\|0 | 1\|0 | 2\|0 | 0\|1 | 1\|1 | 2\|1 | 0\|2 | 1\|2 | 2\|2 |
| Best-guess |  |  |  |  |  |  |  |  |  |  |  |  |
| REF | 129,435 | 81.4 | 90.7 | 98.53 | 1.34 | 0.13 | 24.27 | 52.40 | 23.33 | 0.18 | 1.61 | 98.21 |
| ALT | 129,327 | 81.4 | 90.7 | 98.36 | 1.49 | 0.14 | 23.66 | 52.42 | 23.92 | 0.17 | 1.47 | 98.36 |
| CIS | 129,610 | 81.5 | 90.7 | 98.37 | 1.49 | 0.14 | 23.82 | 52.73 | 23.45 | 0.17 | 1.62 | 98.21 |
| TRANS | 129,148 | 81.3 | 90.6 | 98.52 | 1.34 | 0.13 | 24.11 | 52.10 | 23.79 | 0.18 | 1.46 | 98.36 |
| Probability ≥ 0.90 |  |  |  |  |  |  |  |  |  |  |  |  |
| REF | 43,643 | 95.1 | 97.5 | 97.18 | 2.82 | 0.00 | 3.52 | 93.26 | 3.21 | 0.00 | 3.65 | 96.35 |
| ALT | 43,489 | 95.0 | 97.5 | 96.75 | 3.25 | 0.00 | 3.35 | 93.30 | 3.36 | 0.00 | 3.22 | 96.78 |
| CIS | 43,970 | 95.0 | 97.5 | 96.88 | 3.12 | 0.00 | 3.44 | 93.30 | 3.26 | 0.00 | 3.52 | 96.48 |
| TRANS | 43,145 | 95.1 | 97.6 | 97.10 | 2.90 | 0.00 | 3.42 | 93.28 | 3.30 | 0.00 | 3.32 | 96.68 |
| Probability ≥ 0.98 |  |  |  |  |  |  |  |  |  |  |  |  |
| REF | 10,679 | 98.8 | 99.4 | 99.65 | 0.35 | 0.00 | 1.00 | 98.28 | 0.72 | 0.00 | 0.57 | 99.43 |
| ALT | 10,638 | 98.8 | 99.4 | 99.64 | 0.36 | 0.00 | 0.92 | 98.26 | 0.81 | 0.00 | 0.41 | 99.59 |
| CIS | 10,858 | 98.8 | 99.4 | 99.65 | 0.35 | 0.00 | 0.98 | 98.23 | 0.78 | 0.00 | 0.55 | 99.45 |
| TRANS | 10,463 | 98.8 | 99.4 | 99.64 | 0.36 | 0.00 | 0.94 | 98.31 | 0.75 | 0.00 | 0.43 | 99.57 |

<sup>a</sup> Number of genotypes called for 5,531 SNPs across 26 individuals. Read counts for low-coverage data can be recalled at all variant positions discovered with high-coverage data.

### Index hopping

Index hopping was estimated to be around 1.5% in our dataset. The results of using the method based on presence/absence of each allele, which is more sensitive to index hopping, are shown in Table 8. In the table we show the genotype concordance observed in the real and simulated data. The regression of the genotype concordance for homozygotes on the level of index hopping had very high R-squared (R^2^≥0.99), while R-squared was below 0.05 for heterozygotes. Similarly, the regression of ilr transformations of concordance on the level of index hopping also had a high R-squared when calculated for homozygotes (R^2^≥0.98). In all cases the index hopping level was estimated to be in the range from 1.3% to 1.8%.

**Table 8.** Estimates of index hopping incidence through concordance between low- and high-coverage sequence data in the real and simulated datasets, expressed as percentages or their isometric log-ratios.

|  | Concordance by genotype (%) |  |  |  |  |  |  |  |  | Isometric log-ratios |  |  |  |  |
| --- | --- | --- | --- | --- | --- | --- | --- | --- | --- | --- | --- | --- | --- | --- |
|  | True=0 |  |  | True=1 |  |  | True=2 |  |  | 3 parts<sup>a</sup> |  |  | 2 parts<sup>b</sup> |  |
|  | 0\|0 | 1\|0 | 2\|0 | 0\|1 | 1\|1 | 2\|1 | 0\|2 | 1\|2 | 2\|2 | 0\|0 vs.<br>1\|0, 2\|0 | 1\|1 vs.<br>0\|1, 2\|1 | 2\|2 vs.<br>0\|2, 1\|2 | 0\|0 vs.<br>1\|0 | 2\|2 vs.<br>1\|2 |
| Observed | 98.45 | 1.42 | 0.13 | 24.15 | 52.62 | 23.23 | 0.18 | 1.71 | 98.10 | 4.44 | 0.65 | 4.22 | 3.00 | 2.86 |
| Simulated |  |  |  |  |  |  |  |  |  |  |  |  |  |  |
| 0% | 99.62 | 0.35 | 0.03 | 23.57 | 52.98 | 23.45 | 0.04 | 0.47 | 99.48 | 5.61 | 0.66 | 5.35 | 4.00 | 3.78 |
| 0.1% | 99.53 | 0.44 | 0.03 | 23.59 | 53.52 | 22.89 | 0.08 | 0.52 | 99.40 | 5.47 | 0.68 | 5.06 | 3.83 | 3.72 |
| 0.5% | 99.28 | 0.66 | 0.06 | 23.91 | 53.22 | 22.87 | 0.10 | 0.92 | 98.98 | 5.07 | 0.67 | 4.72 | 3.55 | 3.31 |
| 1% | 98.99 | 0.90 | 0.10 | 23.70 | 53.23 | 23.07 | 0.14 | 1.33 | 98.53 | 4.72 | 0.67 | 4.43 | 3.32 | 3.04 |
| 2% | 98.20 | 1.64 | 0.16 | 23.73 | 52.90 | 23.37 | 0.23 | 2.16 | 97.62 | 4.29 | 0.66 | 4.04 | 2.89 | 2.70 |
| 5% | 96.34 | 3.29 | 0.37 | 23.56 | 53.37 | 23.07 | 0.59 | 4.75 | 94.66 | 3.65 | 0.68 | 3.29 | 2.39 | 2.12 |
| Regression |  |  |  |  |  |  |  |  |  |  |  |  |  |  |
| R<sup>2</sup> | 0.999 | 0.998 | 0.999 | 0.014 | 0.044 | 0.014 | 0.989 | 1.000 | 0.999 | 0.993 | 0.213 | 0.981 | 0.995 | 0.989 |
| Estimate | 1.74 | 1.77 | 1.47 | - | - | - | 1.28 | 1.45 | 1.43 | 1.58 | - | 1.48 | 1.67 | 1.46 |

<sup>a</sup> The 3-part isometric log-ratios take the form $\sqrt{\frac{2}{3}} \ln \frac{(0|0)}{\sqrt{(1|0) \cdot (2|0)}}$
<sup>b</sup> The 2-part isometric log-ratios take the form $\frac{1}{\sqrt{2}} \ln \frac{(0|0)}{(1|0)}$

The results obtained using the concordance variables of best-guess genotypes and genotypes called with probabilities above 0.90 and 0.98, largely supported the results of the presence/absence calling method (data not provided). The results obtained using the concordance variables of best-guess genotypes gave estimates ranging from 1.3% to 1.8% (R^2^≥0.99). The concordance variables of genotypes with probabilities above 0.98 were less sensitive to index hopping and resulted in a lower regression fit and lower or unreliable estimates (1.1% to 1.3%, R^2^=0.96–99, for percentages; 1.4% to 1.7% but R^2^=0.81–0.97 for ilr). The concordance variables of the genotypes with probabilities above 0.90 were in between, with estimates ranging from 1.3% to 1.5% (R^2^≥0.99).

The impact of different levels of index hopping on the genotype concordance is shown in Table 9. Incidences of 1% or 2% of index hopping increased the percentage of incorrect calls from 17.8% to 18.1% or 18.7%, respectively, for best-guess genotypes, from 3.1% to 3.8% or 4.6%, respectively, for genotypes with a probability above 0.90, and from 0.6% to 0.8% or 0.7%, respectively, for genotypes with a probability above 0.98.

**Table 9.** Impact of index hopping level on concordance between low- and high-coverage sequence.

|  | Genotype<br>concordance<br>(%) | Allele<br>concordance<br>(%) | Concordance by genotype (%) |  |  |  |  |  |  |  |  |
| --- | --- | --- | --- | --- | --- | --- | --- | --- | --- | --- | --- |
|  |  |  | True=0 |  |  | True=1 |  |  | True=2 |  |  |
|  |  |  | 0\|0 | 1\|0 | 2\|0 | 0\|1 | 1\|1 | 2\|1 | 0\|2 | 1\|2 | 2\|2 |
| Best-guess |  |  |  |  |  |  |  |  |  |  |  |
| 0% | 82.2 | 91.1 | 99.63 | 0.34 | 0.03 | 23.65 | 52.82 | 23.53 | 0.04 | 0.47 | 99.49 |
| 0.1% | 82.4 | 91.2 | 99.55 | 0.41 | 0.03 | 23.67 | 53.36 | 22.97 | 0.08 | 0.49 | 99.43 |
| 0.5% | 82.2 | 91.1 | 99.30 | 0.64 | 0.06 | 23.97 | 53.08 | 22.96 | 0.10 | 0.89 | 99.01 |
| 1% | 81.9 | 90.9 | 99.03 | 0.87 | 0.10 | 23.77 | 53.09 | 23.14 | 0.14 | 1.28 | 98.58 |
| 2% | 81.3 | 90.6 | 98.25 | 1.59 | 0.16 | 23.79 | 52.78 | 23.43 | 0.23 | 2.07 | 97.70 |
| 5% | 80.1 | 89.9 | 96.45 | 3.17 | 0.37 | 23.63 | 53.23 | 23.14 | 0.59 | 4.58 | 94.82 |
| Probability ≥ 0.90 |  |  |  |  |  |  |  |  |  |  |  |
| 0% | 96.9 | 98.5 | 99.20 | 0.80 | 0.00 | 2.46 | 94.98 | 2.55 | 0.00 | 1.16 | 98.84 |
| 0.1% | 96.7 | 98.3 | 99.01 | 0.99 | 0.00 | 2.59 | 94.70 | 2.71 | 0.00 | 1.26 | 98.74 |
| 0.5% | 96.4 | 98.2 | 98.39 | 1.61 | 0.00 | 2.59 | 94.75 | 2.66 | 0.00 | 1.98 | 98.02 |
| 1% | 96.2 | 98.1 | 97.87 | 2.13 | 0.00 | 2.68 | 94.99 | 2.33 | 0.00 | 2.98 | 97.02 |
| 2% | 95.4 | 97.7 | 96.36 | 3.64 | 0.00 | 2.59 | 94.85 | 2.56 | 0.00 | 4.92 | 95.08 |
| 5% | 93.2 | 96.6 | 92.53 | 7.47 | 0.00 | 2.65 | 94.78 | 2.57 | 0.00 | 10.55 | 89.45 |
| Probability ≥ 0.98 |  |  |  |  |  |  |  |  |  |  |  |
| 0% | 99.4 | 99.7 | 100.00 | 0.00 | 0.00 | 0.53 | 99.03 | 0.44 | 0.00 | 0.00 | 100.00 |
| 0.1% | 99.3 | 99.6 | 100.00 | 0.00 | 0.00 | 0.63 | 98.85 | 0.53 | 0.00 | 0.00 | 100.00 |
| 0.5% | 99.4 | 99.7 | 99.95 | 0.05 | 0.00 | 0.37 | 99.09 | 0.54 | 0.00 | 0.09 | 99.91 |
| 1% | 99.2 | 99.6 | 99.82 | 0.18 | 0.00 | 0.51 | 98.97 | 0.52 | 0.00 | 0.46 | 99.54 |
| 2% | 99.1 | 99.5 | 99.47 | 0.53 | 0.00 | 0.52 | 98.89 | 0.59 | 0.00 | 0.75 | 99.25 |
| 5% | 98.6 | 99.3 | 98.42 | 1.58 | 0.00 | 0.59 | 98.94 | 0.47 | 0.00 | 2.83 | 97.17 |

## Discussion

We quantified the impact of different sources of sequencing errors and biases towards the reference allele on genotype calls derived from low-coverage data. Index hopping and bias towards the reference allele due to alignment had little impact on the genotype calls. However, we found that variant callers can introduce a strong bias towards the reference allele and this has a great impact on genotype calls. This bias is likely to be pipeline specific [6], but we have detected it using one of the most popular tools for variant discovery. The step that causes this bias was designed for the processing of high-coverage data, but introduces a systematic bias when it is applied to low-coverage data. Other unexpected biases may appear when tools that have been designed for use with high-coverage data are used to process low-coverage data. Awareness of these biases allowed us to design a pipeline that gave significantly more accurate genotype calls from low-coverage sequence data than a standard pipeline. In what follows we discuss each of the sources of errors and biases that we have analysed and our proposed new pipeline for variant discovery and joint genotyping, which addresses the most important source of bias.

### Bias towards reference allele due to variant caller

Tools designed for high-coverage sequence data can introduce unexpected biases when used to process low-coverage sequence data. We found this to be the case for the ‘pruning’ step implemented in GATK HaplotypeCaller. During variant discovery it is virtually impossible to distinguish between a sequencing error and a genuine variant. In order to make variant discovery more robust different tools use different strategies to try to identify potential sequencing errors. In the case of GATK HaplotypeCaller, that strategy is the ‘pruning’ step. GATK HaplotypeCaller performs local re-assembly of the reads to generate a list of possible haplotypes in a region by constructing a read-threading graph. Paths of this graph that are supported by a number of reads (kmers) equal or lower than a predefined threshold are considered to probably be sequencing errors and are removed from the graph (pruned). In the next step of the HaplotypeCaller method, each individual read is aligned against each possible haplotype, including the reference, and a likelihood score is calculated for each read-haplotype pair. Then, the likelihood that a read carries each of the alleles at a site is calculated as the product of the likelihoods of all haplotypes that carry that allele. Finally, the allele with the greatest marginal likelihood is called.

While this is a reasonable strategy for high-coverage sequence data, it introduces a huge bias towards the reference allele when used for low-coverage sequence data. This can be intuitively understood with a simple example. Imagine that in any given site with the reference allele ‘A’ and the alternative allele ‘B’ we have only 1 read and that this read carries the alternative allele B. The graph path representing the haplotype with the allele B will be supported by only 1 read and will be pruned out of the graph with the default settings, where at least 2 reads supporting a path are required. This means that the only haplotype that will remain in the graph path is the reference haplotype with allele A. Then, in the next step that same read with the allele B will be paired to all the possible haplotypes. In this case, the only possibility is the reference haplotype with allele A and therefore that read is called as carrying the reference allele A. Thus, instead of the true state with nRef=0 and nAlt=1 we end up with the opposite situation with nRef=1 and nAlt=0. The same bias would arise with a coverage of 3x, if 2 reads carry allele A and 1 read carries allele B. In that case, instead of the true state with nRef=2 and nAlt=1, which indicates a heterozygote, we end up with nRef=3 and nAlt=0, which indicates a reference homozygote. These biased allele read counts are then stored in a gVCF, the file that includes both the variant and non-variant sites and that is used for multi-sample joint genotyping.

The bias in our low-coverage data was so pervasive that it was carried over to their downstream analyses. The bias affected imputation accuracy at population level. We estimated that the individual-wise dosage correlations decreased by an average of 0.10 (0.04 SD; max. 0.20) and the individual-wise percentage of correct best-guess genotypes by 7.5 absolute percentage points (3.8% SD; max. 14.7%) due to this bias (unpublished data). The imputation algorithm that we used for this test calculates genotype probabilities from the allele read counts [21], but the impact of the bias on imputation accuracy could be even greater for imputation algorithms that instead take genotype calls as an input.

#### New pipeline

We propose a new pipeline for variant discovery and genotype calling with low-coverage sequence data. The pipeline that we propose has two steps: (i) variant discovery with the default pruning setting of GATK HaplotypeCaller; and (ii) genotype calling from the aligned reads stored in the BAM files for the variants discovered. This new pipeline gave better genotype and allele concordances than using GATK HaplotypeCaller with disabled pruning.

*- Variant discovery with GATK HaplotypeCaller*: Disabling pruning does not seem an appropriate solution for variant discovery with low-coverage sequencing because this increases the number of potential false positives (Table 2), as well as computational time. The pruning option of GATK HaplotypeCaller makes variant discovery more robust to false positives, but there is a trade-off between specificity and sensitivity. While pruning reduces the ability to discover variants from low-coverage data, this may be overcome by sequencing strategies that target haplotypes from the population instead of individuals (e.g., AlphaSeqOpt; [22,23]) in two ways: (i) sequencing at high coverage of individuals that share large amounts of haplotypes with the population ensures the discovery of many common variants [24]; and (ii) given that the realized coverage at a base position follows a Poisson distribution and, therefore, every individual has greater coverage than the average target coverage in many random positions, many variants can be discovered if a sufficiently large number of individuals are sequenced at low coverage, even if pruning is enabled. For instance, with only 26 individuals sequenced at 2x we discovered 76.3% of the variants discovered with the same individuals at 30x. The gap between variants discovered at low or high coverage would diminish with increasing sample sizes.

*- Genotype calling from aligned reads*: GATK HaplotypeCaller with pruning induces a bias towards the reference allele when used with low-coverage data. This bias is introduced during variant discovery, but manifests itself in the genotype calls because the joint genotyping uses the allele read counts stored in gVCF files produced by the variant caller. This bias can be avoided if we call genotypes based on allele read counts extracted directly from the aligned reads in the BAM files. Tools such as pysam (https://github.com/pysam-developers/pysam) can be used for this purpose.

This pipeline improves scalability. Because gVCF files are created during the variant discovery step, they contain biased allele read counts. In the new pipeline we use regular VCF files to obtain a list of variant positions discovered across the sequenced samples from which we want to extract the raw allele read counts. Using this pipeline it is very easy to add new batches of samples without having to repeat joint genotyping, just by extracting the allele read counts for the new animals and the new variants discovered and then adding them to any pre-existing dataset.

### Bias towards the reference allele due to alignment

With the current pig reference genome *Sscrofa11.1* the bias towards the reference allele due to alignment was very low and its impact on genotype calls was negligible. Our estimates suggest that 1.3% of reads did not align because the reference genome contained the opposite allele to the read allele and this increased the percentage of incorrect best-guess genotype calls by 0.1 absolute percentage points. The reference genome *Sscrofa11.1* was largely constructed using Pacific Biosciences long reads, with a coverage of 65x. This current version of the reference genome provides much better mapping quality than the previous version *Sscrofa10.2* (GenBank accession: GCA_000003025.4). For example, in a 2x sample the percentage of mapped reads increased from 89% in *Sscrofa10.2* to 95% in *Sscrofa11.1*, the percentage of properly paired reads from 77% to 86%, and the percentage of reads with high mapping quality (MAPQ≥40) from 71% to 84%. Here we only considered SNPs but we expect that the alignment bias would have greater impact when using a lower quality reference genome or in regions of high variability and structural complexity, e.g., in presence of multiple indels. The development of alternative-aware alignment algorithms or genome variation graphs [5,25] could alleviate the bias towards the reference genome due to alignment in the near future, but these methods still pose some practical limitations and their use is not generalised yet.

### Index hopping

We estimated the level of index hopping in the 26 samples sequenced in a multiplex at 2x to be 1.5%. This was within the expectation according to Illumina guidelines (<2%; [11]). The impact of index hopping on the percentage of incorrect genotype calls depends on the conservativeness of the calls. For conservative calls the impact was negligible, but for best-guess genotypes the percentage of incorrect calls increased by 0.3 to 0.9 absolute percentage points (1.8% to 5.2% more incorrect calls).

We used a novel empirical method to estimate the level of index hopping. Our method relies on sequencing the same set of samples twice, with multiplexing and without it, so that the level of index hopping in the multiplexed data can be measured against a scale of simulated index hopping levels obtained from a set of index hopping-free data. Previously, Owens et al. [8] proposed a method for testing index hopping that was based on finding heterozygotes with unbalanced read counts for the reference and alternative alleles (e.g., one allele A supported by many reads but the opposite allele B only by one), and then estimating index hopping from the expectation derived from the number of individuals in the multiplex that had many reads supporting that allele B. That method has the advantage that it uses existing data and it does not require the same samples to be sequenced twice. However, that method requires high-coverage data and does not answer how index hopping affects the genotype calls.

Results in our study, together with those of other studies [8,9], reassure us that the high levels of index hopping reported by Sinha et al. [7] are unlikely to occur in most applications if good cleaning protocols are followed to remove excess free-floating indexing primers during library preparation or if unique dual indexed are used [10].

## Conclusion

Index hopping and bias towards the reference allele due to alignment have little impact on downstream genotype calls from low-coverage sequence data, but unexpected biases may arise from pipelines that use tools designed for high-coverage sequence data when used on low-coverage sequence data. The step of ‘pruning’ implemented in GATK HaplotypeCaller is an example of a desirable feature for high-coverage data that introduces a systematic bias when it is applied to low-coverage data. We propose a simple new pipeline to correct this bias. We recommend that users of low-coverage sequencing be very wary of unexpected biases before using tools designed for high-coverage sequencing.

## Declarations

### Ethics approval and consent to participate

The samples used in this study were derived from the routine breeding activities of PIC.

### Consent for publication

Not applicable.

### Availability of data and materials

The datasets generated and analysed in this study are derived from the PIC breeding programme and not publicly available.

### Competing interests

The authors declare that they have no competing interests.

### Funding

The authors acknowledge the financial support from the BBSRC ISPG to The Roslin Institute BBS/E/D/30002275, from Genus plc, Innovate UK (grant 102271), and from grant numbers BB/N015339/1, BB/L020467/1, and BB/M009254/1. M. Johnsson acknowledges financial support from the Swedish Research Council Formas Dnr 2016–01386.

### Authors’ contributions

RRF, AJM, SDR and JHM designed the study; RRF performed the analyses; RRF wrote the first draft; MB, MJ, GG, AJM, SDR, and JHM assisted in the interpretation of the results and provided comments on the manuscript. All authors read and approved the final manuscript.

## Acknowledgements

This work has made use of the resources provided by the Edinburgh Compute and Data Facility (ECDF) (http://www.ecdf.ed.ac.uk/).

## References

1. Li Y, Sidore C, Kang HM, Boehnke M, Abecasis GR. Low-coverage sequencing: Implications for design of complex trait association studies. Genome Res. 2011;21:940–51.

2. Hickey JM. Sequencing millions of animals for genomic selection 2.0. J Anim Breed Genet. 2013;130:331–2.

3. Hickey JM, Gorjanc G, Cleveland MA, Kranis A, Jenko J, Mésázros G, et al. Sequencing Millions of Animals for Genomic Selection 2.0. Proc 10th World Congr Genet Appl Livest Prod WCGALP. Vancouver, BC, Canada; 2014. p. 377.

4. Chen X, Listman JB, Slack FJ, Gelernter J, Zhao H. Biases and Errors on Allele Frequency Estimation and Disease Association Tests of Next-Generation Sequencing of Pooled Samples. Genet Epidemiol. 2012;36:549–60.

5. Li H. Toward better understanding of artifacts in variant calling from high-coverage samples. Bioinformatics. 2014;30:2843–51.

6. Wall JD, Tang LF, Zerbe B, Kvale MN, Kwok P-Y, Schaefer C, et al. Estimating genotype error rates from high-coverage next-generation sequence data. Genome Res. 2014;24:1734–9.

7. Sinha R, Stanley G, Gulati GS, Ezran C, Travaglini KJ, Wei E, et al. Index Switching Causes “Spreading-Of-Signal” Among Multiplexed Samples In Illumina HiSeq 4000 DNA Sequencing. 2017 [cited 2018 Jun 25]; Available from: http://biorxiv.org/lookup/doi/10.1101/125724

8. Owens GL, Todesco M, Drummond EBM, Yeaman S, Rieseberg LH. A novel post hoc method for detecting index switching finds no evidence for increased switching on the Illumina HiSeq X. Mol Ecol Resour. 2018;18:169–75.

9. van der Valk T, Vezzi F, Ormestad M, Dalen L, Guschanski K. Estimating the rate of index hopping on the Illumina HiSeq X platform. 2018 [cited 2018 Jun 25]; Available from: http://biorxiv.org/lookup/doi/10.1101/179028

10. Costello M, Fleharty M, Abreu J, Farjoun Y, Ferriera S, Holmes L, et al. Characterization and remediation of sample index swaps by non-redundant dual indexing on massively parallel sequencing platforms. BMC Genomics. 2018;19.

11. Illumina Inc. Effects of Index Misassignment on Multiplexing and Downstream Analysis [Internet]. 2017. Available from: https://www.illumina.com/content/dam/illuminamarketing/documents/products/whitepapers/index-hopping-white-paper-770-2017-004.pdf

12. Brandt DYC, Aguiar VRC, Bitarello BD, Nunes K, Goudet J, Meyer D. Mapping Bias Overestimates Reference Allele Frequencies at the *HLA* Genes in the 1000 Genomes Project Phase I Data. G3amp58 GenesGenomesGenetics. 2015;5:931–41.

13. Poplin R, Ruano-Rubio V, DePristo MA, Fennell TJ, Carneiro MO, Van der Auwera GA, et al. Scaling accurate genetic variant discovery to tens of thousands of samples. 2017 [cited 2018 Jun 28]; Available from: http://biorxiv.org/lookup/doi/10.1101/201178

14. DePristo MA, Banks E, Poplin R, Garimella KV, Maguire JR, Hartl C, et al. A framework for variation discovery and genotyping using next-generation DNA sequencing data. Nat Genet. 2011;43:491–8.

15. Bolger AM, Lohse M, Usadel B. Trimmomatic: a flexible trimmer for Illumina sequence data. Bioinformatics. 2014;30:2114–20.

16. Li H, Durbin R. Fast and accurate short read alignment with Burrows-Wheeler transform. Bioinforma Oxf Engl. 2009;25:1754–60.

17. Danecek P, Auton A, Abecasis G, Albers CA, Banks E, DePristo MA, et al. The variant call format and VCFtools. Bioinformatics. 2011;27:2156–8.

18. Aitchison J. The statistical analysis of compositional data. London; New York: Chapman and Hall; 1986.

19. Egozcue JJ, Pawlowsky-Glahn V, Mateu-Figueras G, Barceló-Vidal C. Isometric Logratio Transformations for Compositional Data Analysis. Math Geol. 2003;35:279–300.

20. Ros-Freixedes R, Estany J. On the Compositional Analysis of Fatty Acids in Pork. J Agric Biol Environ Stat. 2014;19:136–55.

21. Whalen A, Ros-Freixedes R, Wilson DL, Gorjanc G, Hickey JM. Hybrid peeling for fast and accurate calling, phasing, and imputation with sequence data of any coverage in pedigrees. 2017 [cited 2018 Jun 28]; Available from: http://biorxiv.org/lookup/doi/10.1101/228999

22. Gonen S, Ros-Freixedes R, Battagin M, Gorjanc G, Hickey JM. A method for the allocation of sequencing resources in genotyped livestock populations. Genet Sel Evol. 2017;49:47.

23. Ros-Freixedes R, Gonen S, Gorjanc G, Hickey JM. A method for allocating low-coverage sequencing resources by targeting haplotypes rather than individuals. Genet Sel Evol. 2017;49:78.

24. Xu C, Wu K, Zhang J-G, Shen H, Deng H-W. Low-, high-coverage, and two-stage DNA sequencing in the design of the genetic association study. Genet Epidemiol. 2017;41:187–97.

25. Paten B, Novak AM, Eizenga JM, Garrison E. Genome graphs and the evolution of genome inference. Genome Res. 2017;27:665–76.

